# Genome-Wide Uncertainty-Moderated Extraction of Signal Annotations from Multi-Sample Functional Genomics Data

**DOI:** 10.1101/2025.02.05.636702

**Authors:** Nolan H. Hamilton, Yu-Chen E. Huang, Benjamin D. McMichael, Michael I. Love, Terrence S. Furey

**Affiliations:** Department of Genetics, University of North Carolina at Chapel Hill, Chapel Hill, NC, USA; Department of Biology, University of North Carolina at Chapel Hill, Chapel Hill, NC, USA; Department of Biostatistics, University of North Carolina at Chapel Hill, Chapel Hill, NC, USA; Curriculum in Bioinformatics and Computational Biology, University of North Carolina at Chapel Hill, Chapel Hill, NC, USA

**Keywords:** functional genomics, genomic signal processing, cohort-level estimation, epigenomics

## Abstract

Multi-sample functional genomics experiments should reveal reproducible regulatory activity, but locus- and sample-specific noise can obscure biological signals in sequencing data. We introduce `Consenrich` for genome-wide estimation of epigenomic signals across multiple samples. To encourage robustness and sensitivity, the state-space model underlying `Consenrich` accounts for positional observation variances to determine shrinkage toward predictions from a smooth process model over genomic coordinates. We first apply `Consenrich` to ATAC-seq and ChIP-seq datasets and demonstrate its ability for robust signal recovery. We then utilize `Consenrich` upstream of a class-imbalanced differential accessibility analysis in an Alzheimer’s cohort of twenty samples and show that it improves the breadth of relevant biological insights. Software is available at https://github.com/nolan-h-hamilton/Consenrich.

## 1 Background

High-throughput functional genomics assays measure diverse molecular phenotypes across the genome, including chromatin accessibility, protein-DNA interactions, and chromatin-associated modifications. Examples include the assay for transposase-accessible chromatin with sequencing (ATAC-seq), DNase I hypersensitive-site sequencing (DNase-seq), chromatin immunoprecipitation sequencing (ChIP-seq), and cleavage under targets and release using nuclease (CUT&RUN). These assays generate sequence reads that are aligned to the genome and summarized as regions with enriched read coverage or signal, commonly referred to as *peaks* [1–3]. These peak annotations can then serve as candidate regulatory elements for downstream analysis, including tests for differential regulatory activity between phenotypically distinct groups of samples. To perform these analyses over discrete regions, a consensus set of peak annotations is defined across all samples. Two sources of noise arise repeatedly in functional genomics data that make consensus peak calling challenging. First, samples can differ in their overall signal quality, such as from technical variation arising when running the assay or from biological variation due to factors such as the quality of the sample material. Second, noise profiles can vary locally across the genome within each sample or generally across samples. For instance, a sample with strong overall signal quality may display poor signal quality at particular loci. Likewise, features of local DNA sequences, such as guanine-cytosine (GC) content, can affect the level and quality of signals generated by an assay. Thus, multi-sample signal estimation requires accounting for both sample-level and locus-level variation in data quality.

Many established peak callers, including Model-based Analysis of ChIP-Seq (`MACS`), `F-seq`, `HMMRATAC`, and `LanceOTron` [3–6], are designed primarily to identify enriched regions in individual samples or datasets. To generate multi-sample consensus peaks using these methods, one strategy is to pool reads across samples and call peaks on the aggregate alignment file [7, 8]. Although simple and computationally convenient, pooling alignments precludes hierarchical treatment of samples based on quality and makes the final signal estimate depend directly on the aggregate read mixture. Several strategies for aggregating sample-level measurements are available through tools like `MACS cmbreps` [3, 9]. However, these approaches still collapse sample information before peak calling and do not explicitly model sample-level quality, sample-specific artifacts, or uncertainty in the consensus signal. In `MACS` workflows, local background signal estimates may also be combined by naive aggregation across multiple scales (1 kilobase (kb) and 10 kb), but these constant scales are arbitrary and do not appeal to concrete biological reasoning or trends observed in the data.

Another popular strategy determines consensus enrichment regions by counting overlaps in each individual sample’s pre-called peaks. The overlapping peaks are then merged to create a consensus annotation using heuristic overlap or recurrence criteria [10–14]. These post hoc approaches make consensus peak annotations dependent on discrete sample-level peak calls, so modest but reproducible enrichment can be missed if it fails to cross individual-sample detection thresholds. Conversely, noisy sample-specific peaks can distort consensus peak boundaries or introduce peaks with weak cohort-level support. Choosing overlap or recurrence criteria is also difficult, particularly in heterogeneous or imbalanced case-control studies where biologically relevant signal may be present in only a subset of samples. Despite these limitations, pooling and post hoc merging remain common practice for consensus peak calling [15–18]. Related tools developed primarily for ChIP-seq data analysis, such as `DiffBind` and `csaw`, are more specifically aimed at differential analyses across sets of phenotypically distinct samples [10, 19]. `DiffBind` typically begins from sample-level peak annotations, constructs a consensus peak annotation with an overlap rule, and then generates a count matrix for input to differential analysis methods like `DESeq2` [20]. `csaw` instead performs differential analyses directly in sliding windows across the genome without relying on a discrete peak annotation.

`Consenrich` is a state-space estimator for multi-sample functional genomics datasets with a loci- and sample-specific observation model to address heterogeneous data quality and a prior process model for transitions between successive genomic intervals. This allows the estimator to favor lower-variance observations while shrinking toward a smooth propagated prediction. Empirically, we evaluate `Consenrich` on ATAC-seq, DNase-seq, and ChIP-seq datasets, focusing on robust recovery of epigenomic signals and determination of consensus peaks for differential analyses.

## 2 Results

### 2.1 Conceptual overview of the Consenrich methodology

`Consenrich` treats the consensus epigenomic signal as an unobserved sequence along the genome. A visual representation of the `Consenrich` method is presented in Figure 1 (see Section 5 for full details). `Consenrich` estimates signal over a partitioned genome consisting of *n* fixed-width intervals (e.g., 50 bp). Observed data over these genomic intervals is a matrix of transformed sample-level signals across the *m* samples in the cohort. At each genomic interval, a smooth linear *process model* gives an *a priori* prediction for interval *i* from the estimate at interval *i* − 1 during the *forward* or *filtering* pass. The *correction* step is then carried out through the *observation model*, which compares the predicted signal to the observed data at interval *i* and applies a weighted correction to the state estimate. This weighted correction is controlled by the *gain*, which arises from the relative uncertainty in the *a priori* prediction and the observed data [21]. Intuitively, when the observations are unreliable relative to the prediction, the gain is low and the subsequent correction is minimal. Conversely, when the observations are reliable the gain is high and the correction is larger. The assignment of uncertainty to the process and observation models is discussed in Section 5. Importantly, uncertainty in the estimates is propagated and updated through the same sequential prediction-correction procedure such that positional covariances are captured and bin-level uncertainty is implied by the model.

**Fig. 1:**
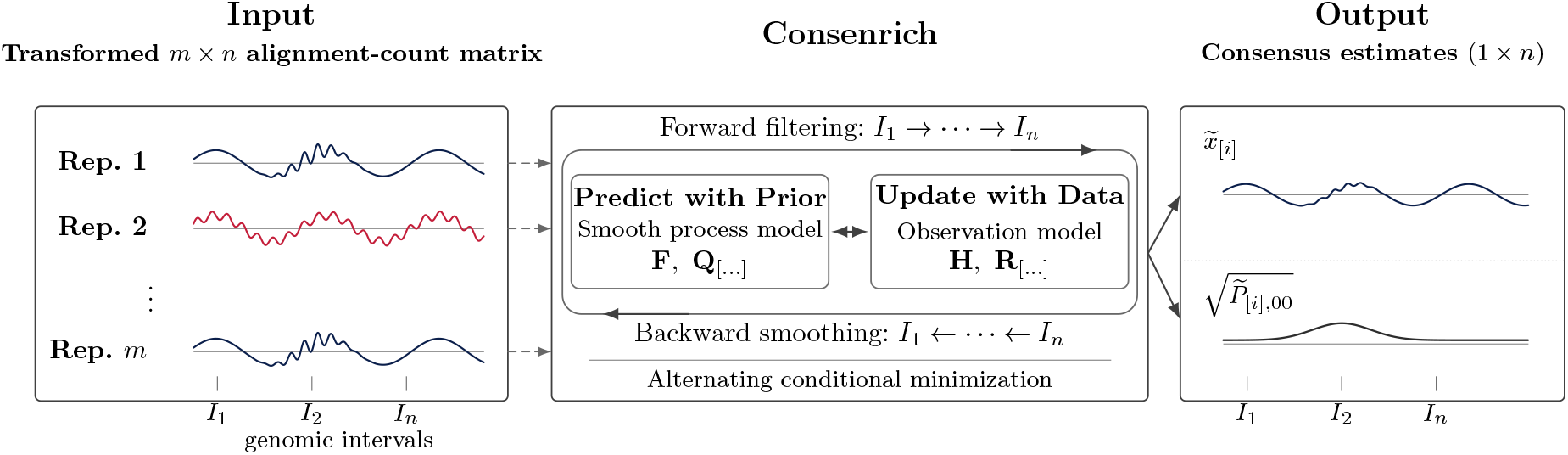
Conceptual schematic. `Consenrich` produces genome-wide signal and uncertainty annotations given multiple sequence alignments. To encourage sufficient robustness and sensitivity, the state-space model underlying `Consenrich` utilizes an observation model {**H**^*m*×2^, **R**^*m* ×*m*^ } with sample- and locus-specific variance to determine shrinkage toward predictions suggested by a smooth, adaptive process model, {**F**^2×2^, **Q**^2×2^}. `Consenrich` fits the corresponding state-space model and nuisance terms on an augmented Kalman filter-smoother loss. Here, 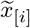 denotes the final state estimate and 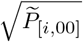 denotes the corresponding model-implied standard deviation. In this example, noisy genomic intervals are represented by interval *I*_2_, where all samples experience instability. Noisy samples are represented by Rep. 2 which is unstable over all intervals.

### 2.2 Robust signal estimation in noisy multi-sample data

A key challenge in combining data across samples is accounting for varying data quality. Data from noisy, low-quality samples can dilute valuable information contained in other samples with less noisy, high-quality data. This challenge is exacerbated in high-throughput sequencing data, where noise processes affecting data across the genome are nonstationary due to region-dependent factors. We first evaluated the estimation performance of `Consenrich` using mixed-quality chromatin accessibility (ATAC-seq) data from a set of lymphoblastoid cell lines with intentionally contaminated synthetic noisy samples.

We used the TSS enrichment score (TSSe), a signal-to-background quality metric defined by ENCODE for evaluation of ATAC-seq data, to rank and define the so-called *Best5* and *Worst5* lymphoblastoid samples from a set of fifty-six total samples [22]. The *Best5* and *Worst5* samples are collectively referred to as the heterogeneous dataset, *Het10*. Additionally, ten synthetic noisy samples, which are comparatively devoid of signal (*Noisy10* ; TSSe ≈ 3.5), were generated and used in evaluations to mimic the effects of low-quality samples in real-world experiments. The *Het10 + Noisy10* dataset is the union of the *Het10* and *Noisy10* samples (Supplementary Section B.1).

For benchmarking, we included two `MACS3 cmbreps` signal quantification summaries, which we refer to as `MACS3 FE (cmbreps)` and `MACS3 logFE (cmbreps)`. The fold-enrichment (FE) uses `macs3 bdgcmp -m FE` to compute a genome-wide enrichment estimate akin to ENCODE’s signal quantification pipelines for ATAC-seq and ChIP-seq experiments [9, 23, 24]. The log fold-enrichment summary uses `macs3 bdgcmp -m logFE` fold-enrichment estimates from the same `MACS3 cmbreps` workflow but on log_10_-scale. A `TSSe-weighted` least squares estimator was also included as a comparator to contrast `Consenrich`’s own sample- and locus-specific perspective with a simpler sample-level weighting approach. Reference feature sets were constructed as putative positive and negative controls for signal enrichment (Section 5.7).

For *Accessible* regions, we used a set of 9,420 regions corresponding to ubiquitous DNase I peak clusters confirmed in multiple cell-line experiments. We likewise compiled 2,392 *Inaccessible* heterochromatic regions from H3K9me3 experiments in the Human Heterochromatin Database (Section 5.7) [25]. Figure 2 demonstrates qualitative estimator behavior over a 25 kb locus.

**Fig. 2:**
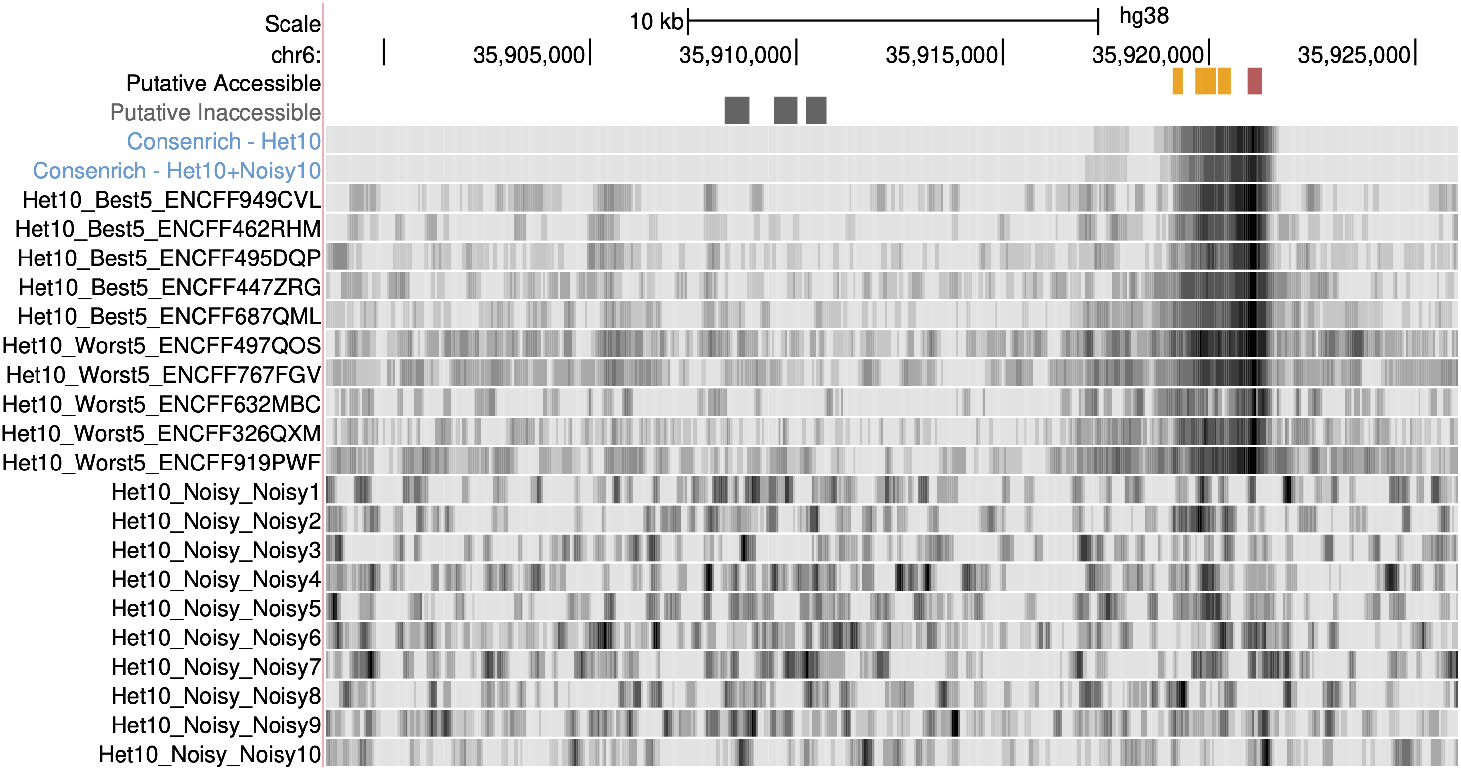
Signal quantification from noisy, multi-sample functional genomics data. Genome-browser signal tracks for `Consenrich` estimates in the *Het10* and *Het10 + Noisy10* ATAC-seq input datasets are shown using the same visualization settings. In each track, color intensity reflects signal amplitude, with darker colors corresponding to higher positive enrichment. At this exemplar locus, `Consenrich` localizes signal near putative accessible annotations, and the *Het10* and *Het10 + Noisy10* tracks retain similar structure.

The first experiments addressed each method’s ability to (i) separate enriched *Accessible* regions or depleted *Inaccessible* regions relative to a null background signal and, (ii) maintain that separation as sample quality degrades. To this end, we computed enrichment statistics, *Z*_boot_, that quantify the differences in the average observed signal from the null distribution. Null distributions were approximated by calculating average signal values over matched-null feature sets generated with `nullranges::bootRanges` (Section 5.6). The underlying null hypothesis is that the observed average over the reference feature set is no stronger than what would be expected for randomly sampled genomic regions with similar local clustering, sizes, and gene densities. For *Accessible* regions, we computed *Z*_boot_ = (*µ*_obs_ − *µ*_null_)*/*(*σ*_null_), where *σ*_null_ is the sample standard deviation of the bootstrapped null distribution. For *Inaccessible* regions, the sign is reversed so that larger values indicate stronger depletion.

In Table 1, all methods cleanly separate *Accessible* regions in the uncontaminated Het10 dataset, with `Consenrich` showing the strongest enrichment in *Accessible* regions. Similarly, `Consenrich` shows the greatest depletion in signal in *Inaccessible* regions. Whereas the comparators show a marked drop in separation after noisy samples are added, `Consenrich` is less affected and shows stronger separation between *Accessible* and *Inaccessible* regions in this scenario.

**Table 1:**
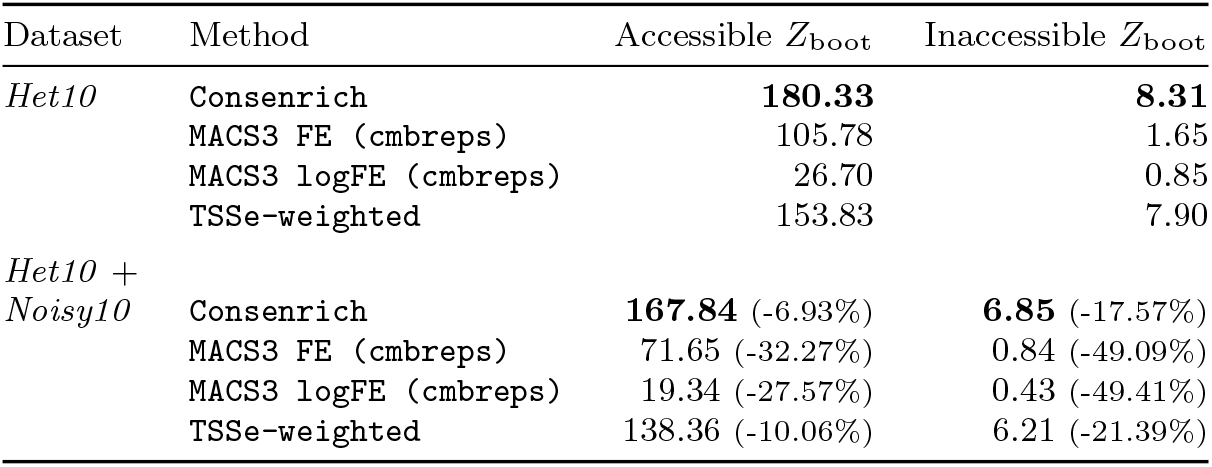
Accessible enrichment and Inaccessible depletion in Het10 and Het10 + Noisy10 datasets. Entries report bootstrapped *z*-statistics relative to matched *Null* feature sets generated with `nullranges::bootRanges`. For the putative Accessible features, enrichment is *Z*_boot_ = (*µ*_obs_ − *µ*_null_)*/*(*σ*_null_). For putative Inaccessible features, depletion is *Z*_boot_ = (*µ*_null_ − *µ*_obs_)*/*(*σ*_null_), so larger values indicate lower signal. The percent differentials in *Het10* + *Noisy10* rows report relative drop from the matching *Het10* value. `Consenrich` shows strong separation from the empirical null distributions for both Accessible and Inaccessible regions and experiences the least degradation in performance with noisy samples added.

| Dataset | Method | Accessible $Z_{\text{boot}}$ | Inaccessible $Z_{\text{boot}}$ |
| --- | --- | --- | --- |
| *Het10* | **Consenrich** | **180.33** | **8.31** |
|  | MACS3 FE (cmbreps) | 105.78 | 1.65 |
|  | MACS3 logFE (cmbreps) | 26.70 | 0.85 |
|  | TSSe-weighted | 153.83 | 7.90 |
| *Het10 + Noisy10* | **Consenrich** | **167.84** (-6.93%) | **6.85** (-17.57%) |
|  | MACS3 FE (cmbreps) | 71.65 (-32.27%) | 0.84 (-49.09%) |
|  | MACS3 logFE (cmbreps) | 19.34 (-27.57%) | 0.43 (-49.41%) |
|  | TSSe-weighted | 138.36 (-10.06%) | 6.21 (-21.39%) |

In addition to the raw separation measure *Z*_boot_, it is also important to consider resilience of *signal structure*: Even if a method maintains strong separation of enriched regions from the null distribution, it may fail to preserve the local shapes of signal within those regions when exposed to genomic artifacts or noisy samples. Mean-dependent effects over the dynamic range of signal values should also be accounted for, since noise profiles can vary in stronger versus weaker signal regions, and genome-wide summary statistics can become dominated by common but uninteresting low-signal blocks. To this end, we evaluated the preservation of local signal shapes within sampled genomic blocks. For each method, we sampled 10 kb blocks and stratified Spearman correlations between *Het10* and *Het10* + *Noisy10* by eight signal quantiles. The resulting median block Spearman correlations within each signal level are shown in Figure 3, where `Consenrich` maintained the highest block Spearman correlations across signal levels despite the addition of noisy samples.

**Fig. 3:**
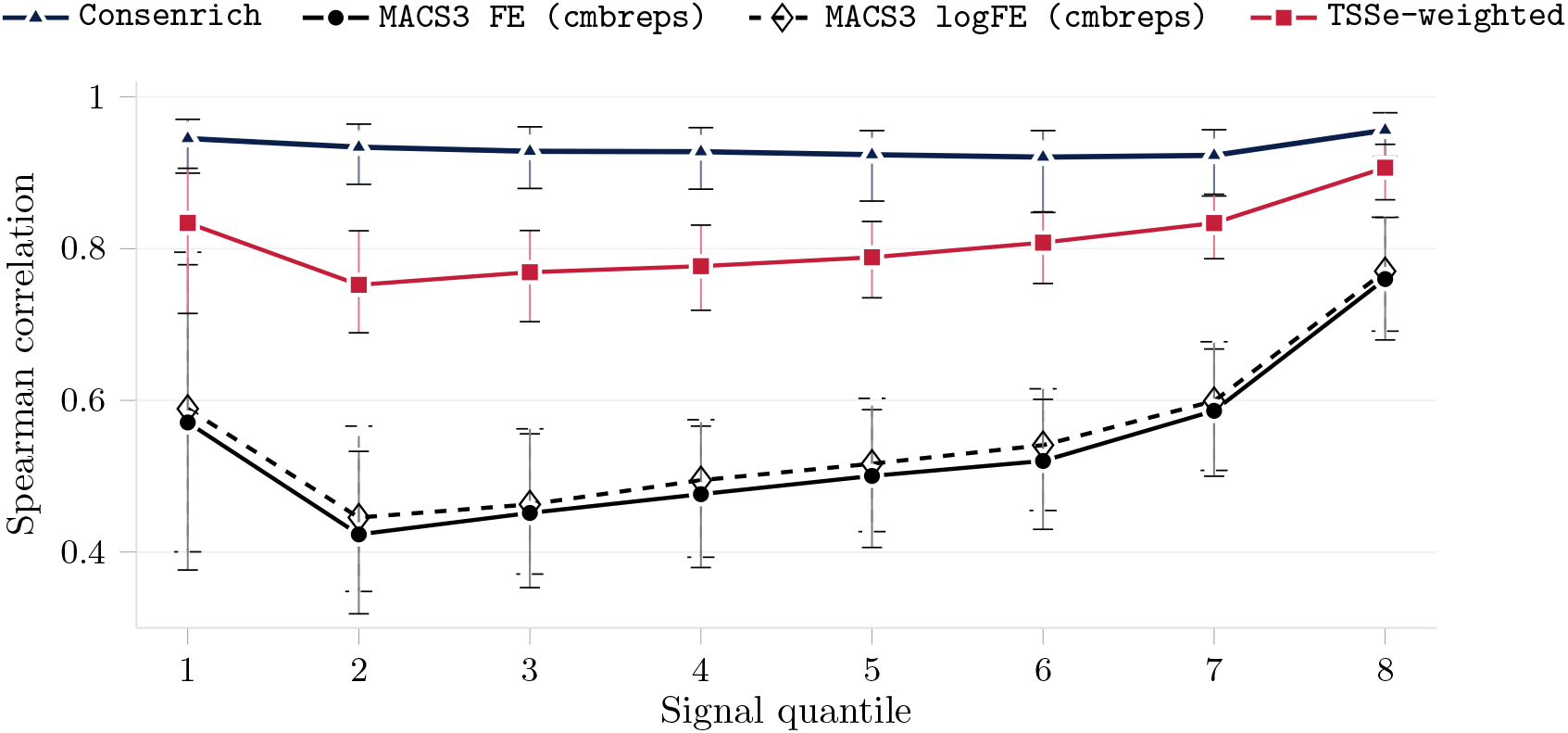
Consenrich preserves local signal rank structure in signal-bearing blocks under added noisy replicates. Genomic blocks with defined within-block Spearman correlations were stratified into 8 within-method signal bins using the *Het10* block mean signal. Points show median Spearman correlation between *Het10* and *Het10 + Noisy10* binned signal values across matched 10 kb genomic blocks, with interquartile ranges shown as vertical bars. Flat blocks have undefined Spearman correlations and are omitted rather than imputed.

As described previously, during the correction step, `Consenrich` calculates a *gain* matrix from the propagated state covariance and the estimated sample-by-interval observation covariance to determine the extent to which each sample contributes to the corrected signal at a given interval. In the *Het10* + *Noisy10* experiment, group-average gains roughly follow the TSSe grouping, with *Best5* samples contributing most on average, *Worst5* samples intermediate on average, and *Noisy10* samples contributing the least (Figure 4). Agreement with this scalar quality-control metric reflects Consenrich’s sample-specific perspective on the varying credibility of replicate data. The within-sample standard deviation around the plotted genome-wide averages then reflects Consenrich’s region-specific perspective on variation in data quality. The gain summaries also showed substantial interval-level spread, illustrating region-specific variation in process and observation uncertainty beyond fixed scalar values. Reporting gains as mean *×*10^3^, *Best5* samples had average gain 15.71 *±* 3.59 over samples, *Worst5* samples had 13.65 *±* 4.84, and *Noisy10* samples had 4.574 *±* 0.007. The mean within-sample gain standard deviation was 2.44 for *Best5*, 2.07 for *Worst5*, and 1.06 for *Noisy10*.

**Fig. 4:**
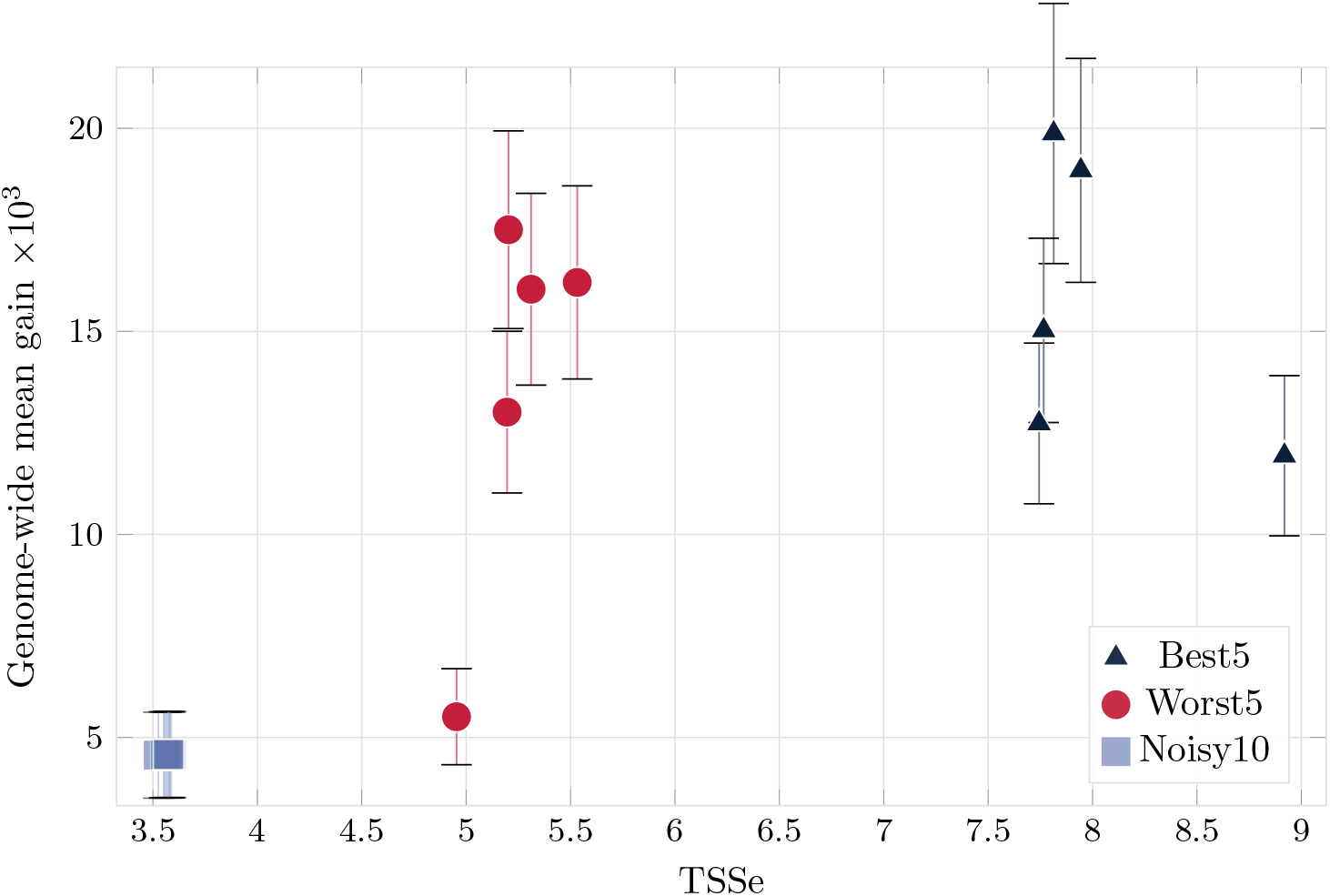
Sample gains versus scalar, enrichment-based quality scores. For each sample in the *Het10 + Noisy10* ATAC-seq experiment, TSSe is plotted against its genome-wide mean gain. The synthetic *Noisy10* replicates cluster strongly at low TSSe and low mean gain, whereas samples with highest TSSe generally receive larger gains. Vertical bars show within-sample standard deviations across genomic intervals, capturing locus-specific variation in each sample’s contribution to the state update. This directional agreement with a model-external quality score is consistent with reduced influence of low-quality samples, while the interval-level spread shows that sample contributions can vary substantially across the genome.

We next evaluated two ChIP-seq datasets with input controls. The input control is meant to help account for non-random signal due to technical or biological noise. Several peak calling methods have been developed for individual ChIP-control sample pairs but methods for the multi-sample case are less established [3, 4, 9]. First we focused on RNA polymerase II subunit A (POLR2A) binding ChIP-seq data from six colon tissue samples. POLR2A binding is expected to be highly enriched near promoter regions of actively transcribed genes. We therefore defined a working positive control set of *Actively Bound* POLR2A loci that overlapped GENCODE-annotated TSS for genes expressed in colon cells [26]. We likewise defined a putative negative control set of *Inaccessible* loci supported by consensus peaks from distinct ENCODE H3K9me3 histone modification ChIP-seq experiments, where this histone modification is associated with inaccessible, inactive chromatin (Section 5.7). Over these regions, we expect that average POLR2A binding signal should be enriched in the *Actively Bound* set and depleted in the *Inaccessible* set. We generated 1,000 *Null* genomic feature sets with`nullranges::bootRanges` (Section 5.6) to approximate a null distribution for testing. As shown in Table 2, `Consenrich` demonstrates substantial enrichment at *Actively Bound* regions and a near-null *Z*_boot_ of 0.12 for putative *Inaccessible* regions. This supports the expected POLR2A-specific pattern in this experiment, while `MACS3 FE (cmbreps)` retains the largest positive-reference statistic.

**Table 2:** ChIP-seq enrichment and depletion summaries. Entries report bootstrapped *z*-statistics for matched reference and null feature sets. For the putative Actively Bound features, larger values indicate stronger enrichment versus the null distribution. For the putative Inaccessible features, larger values indicate stronger depletion versus the null distribution.

| Dataset | Method | Actively Bound $Z_{\text{boot}}$ | Inaccessible $Z_{\text{boot}}$ |
| --- | --- | --- | --- |
| *POLR2A* | **Consenrich** | 35.73 | **0.12** |
|  | MACS3 FE (cmbreps) | **196.78** | -3.19 |
|  | MACS3 logFE (cmbreps) | 15.89 | -0.13 |
| *H3K4me1* | **Consenrich** | **57.70** | **19.65** |
|  | MACS3 FE (cmbreps) | 17.92 | 2.22 |
|  | MACS3 logFE (cmbreps) | 7.37 | 1.01 |

ChIP-seq experiments targeting transcription factor binding and histone modifications are often labeled as ‘narrow’ or ‘broad’ depending on the genomic size of the targeted signal. POLR2A binding signal is considered narrow, with punctate enrichment over bound genomic regions. To also consider broad marks, we evaluated H3K4me1 histone modification datasets with respect to a positive reference set (Section 5.7) defined by consensus peaks in distinct H3K4me1 ENCODE experiments in cell lines. We used the same H3K9me3-based putative inaccessible regions as a working negative reference set. As shown in Table 2, `Consenrich` clearly shows the highest enrichment in consensus H3K4me1 positive regions and the greatest depletion in the H3K9me3 negative regions.

### 2.3 Comprehensive consensus peak calling and differential analyses in complex human disease

We next employed `Consenrich` in an Alzheimer’s disease (AD) case-control differential accessibility analysis. The chromatin accessibility profiles were generated by DNase-seq from dorsolateral prefrontal cortex (DLPFC) tissues of *N*_donor_ = 20 female donors over 75 years of age. Of these donors, *N*_AD_ = 5 had been diagnosed with Alzheimer’s disease (AD), whereas the remaining *N*_nonAD_ = 15 donors did not have AD (non-AD). This 3:1 class imbalance was intended to represent challenging experimental settings with uneven case-control structures [27–30]. Identification of candidate consensus peaks was performed using `Consenrich` and several alternative methods. All consensus peak-calling methods were applied to the full *N*_donor_ = 20 cohort without reference to disease status. Compared with merging peaks identified separately from each condition, this approach has been found to mitigate false positives caused by implicit data snooping [31].

The software implementation of `Consenrich` includes the `ROCCO` [32] peak caller for genome-wide discrete annotation of enriched regions. For `Consenrich+ROCCO`, `ROCCO` was run on the `Consenrich` genome-wide signal estimate to call candidate regions. Standalone `ROCCO`, run directly on the input alignments, is also included as a comparator. We also evaluated `Genrich`, which has been successfully applied to data from several experimental modalities, including ChIP-seq, DNase-seq, and ATAC-seq [33–38]. `Genrich` computes position-wise enrichment *p*-values separately for each sample and combines them across samples using Fisher’s method [39, 40] before calling consensus-enriched candidate regions. Peak calling is additionally controlled by several `Genrich`-specific thresholds, such as the total area under the − log_10_(*p*) significance curve, but these were not incorporated in analyses here. `DiffBind` [10] was also tested. As input, we supplied all 20 samples’ filtered BAM alignments and `MACS3`-based narrowPeak files from ENCODE. `DiffBind` constructed a cohort-wide consensus peak set without reference to disease status by retaining regions supported by at least two samples (`minOverlap=2`) or at least three samples (`minOverlap=3`). Read counts were then obtained from the BAM files in `DiffBind`-defined consensus regions. Finally, because differential accessibility can also be analyzed without first committing to a called peak set, we included a sliding-window method, `csaw` [19]. Unlike the peak-based consensus methods above, `csaw` tiles the genome and merges nearby significant windows after statistical testing. Results should therefore be interpreted in light of this window-based testing universe, which differs from the method-specific consensus regions evaluated by the peak-based methods and limits direct comparability. Note that `csaw` and `DiffBind` focus differential analysis specifically, whereas `Consenrich` is better viewed upstream of differential testing as a generic enrichment estimator whose output can be used for multiple downstream analyses.

With the exception of `csaw`, which directly applies a window-based differential analysis, we performed differential analysis on consensus peak sets using `DESeq2` (Section 5.9). Both explicit covariates, such as donor age, and latent factors estimated from negative-control regions were included in the final `DESeq2` design formula. For peak-based methods, we refer to regions passing | log_2_ FC| *>* 0.25 and FDR-corrected *p*_adj_ *<* 0.05 as differentially accessible regions (DARs). For `csaw`, DARs are significant merged windows returned by `csaw`. In Table 3, we list the number of candidate regions for peak-based methods and merged windows for `csaw`, along with the number found to be differentially accessible with respect to Alzheimer’s disease status.

**Table 3:** For each method listed in the leftmost column, the table reports the total number of candidate regions, the number and percentage of differentially accessible regions (DARs, AD vs non-AD), and the mean and standard deviation of DAR size.

| Method | Candidate Regions | Significant DARs, $N_{\text{DAR}}$ (%) | DAR Size $\hat{\mu}$ (bp) | DAR Size $\hat{\sigma}$ (bp) |
| --- | --- | --- | --- | --- |
| Consenrich+ROCCO | 470,601 | 11,418 (2.4%) | 428.55 | 278.59 |
| ROCCO | 232,087 | 6,043 (2.6%) | 561.96 | 388.48 |
| Genrich | 749,350 | 1,508 (0.2%) | 821.27 | 689.42 |
| `csaw` (5 kb background) | 631,087 | 3,239 (0.5%) | 825.47 | 230.30 |
| `csaw` (10 kb background) | 618,573 | 2,424 (0.4%) | 834.26 | 222.59 |
| DiffBind (`minOverlap=2`) | 148,066 | 5,799 (3.9%) | 588.55 | 211.85 |
| DiffBind (`minOverlap=3`) | 107,651 | 5,929 (5.5%) | 569.91 | 188.07 |

Although downstream multiple-testing correction should ideally control false discoveries across candidate regions at the nominal level, we emphasize that an increased number of significant differential results does not necessarily establish improved detection of informative, disease-relevant regions. We therefore asked whether genes linked to `Consenrich+ROCCO` DARs reflected AD-relevant regulatory programs. Gene-region enrichment testing using Gene Ontology (GO)-defined gene sets [41] with genes assigned to the 11,418 `Consenrich+ROCCO` DARs highlighted neuronal development, neurite growth, cytoskeletal regulation, and synaptic biology. Figure 5 depicts GO enrichment for genes assigned to all `Consenrich+ROCCO` DARs in the left panel and to the 3,979 peak-exclusive `Consenrich+ROCCO` DARs after excluding peak overlaps with ROCCO, Genrich, `csaw`, and DiffBind in the right panel. The strongest raw Biological Process term was axonogenesis, supported by 251 unique genes. Other highly ranked terms included regulation of neuron projection development, regulation of nervous system development, small GTPase-mediated signal transduction, dendrite development, regulation of synapse structure or activity, regulation of neurogenesis, regulation of synapse organization, and gliogenesis. This pattern is concordant with a single-cell transcriptomic study of ROSMAP donors that reported AD-associated gene-expression changes in several brain cell types and identified gene-trait module M3, which overlapped cognition-associated genes and was enriched for terms including axonogenesis, small GTPase mediated signal transduction, and positive regulation of nervous system development [42].

**Fig. 5:**
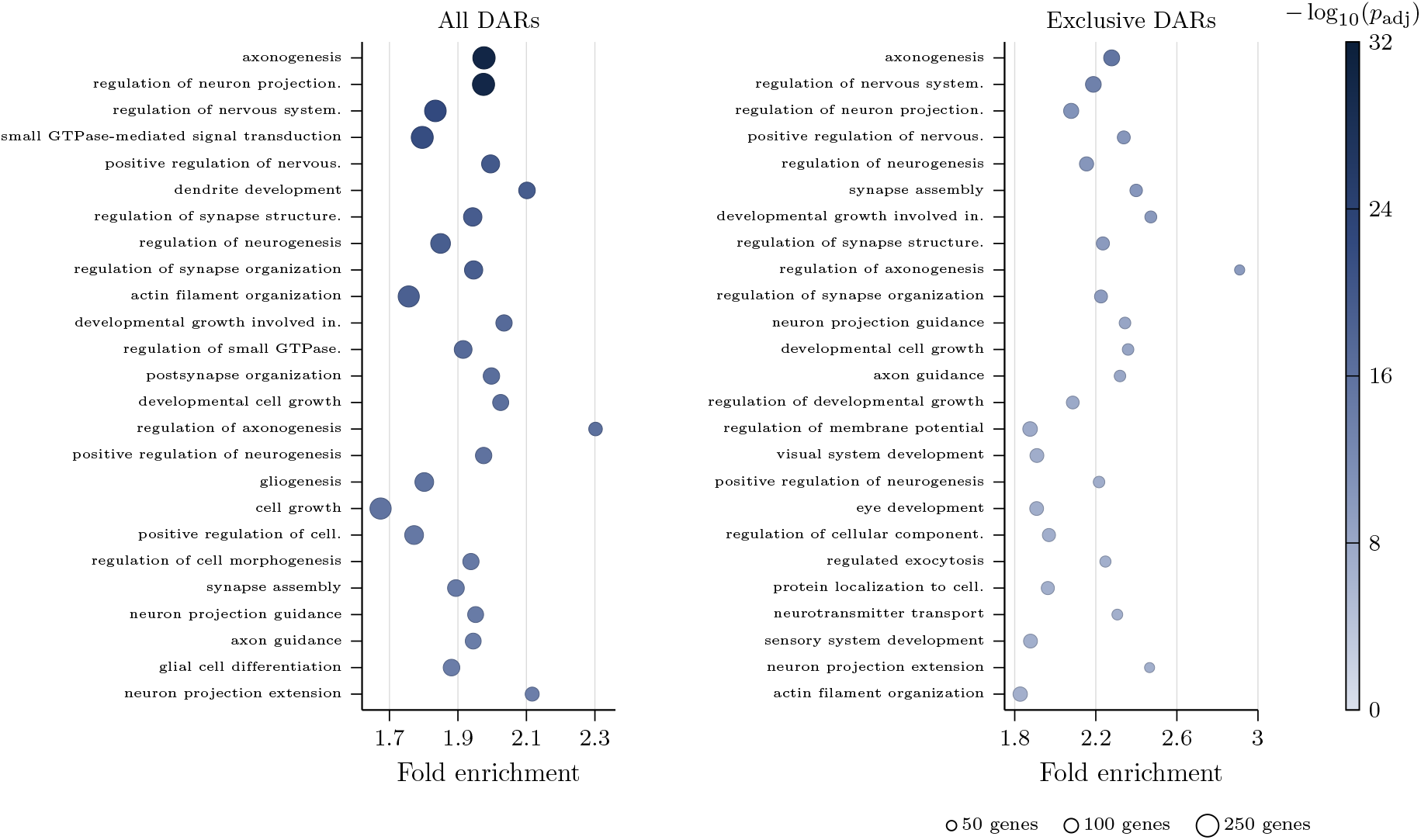
Gene Ontology Biological Process enrichment for genes assigned to Consenrich+ROCCO DARs. The left panel shows terms from all `Consenrich+ROCCO` DARs. The right panel shows terms from the 3,979 peak-exclusive `Consenrich+ROCCO` DARs with no peak overlap with ROCCO, Genrich, `csaw` (5/10 kb), or DiffBind (minOverlap=2/3) results. Points show the 25 raw `enrichGO` terms with smallest adjusted *p*-values.

### 2.4 DAR sets overlap AD risk loci and brain tissue expression quantitative trait loci

To support the relevance of putatively altered regulatory regions in Alzheimer’s disease, we compared DARs to expression quantitative trait loci (eQTLs) identified in ROSMAP dorsolateral prefrontal cortex samples and to genomic risk loci associated with AD. Prior work has shown enrichment of eQTLs in differentially accessible regions in the same tissue, and eQTLs enriched in DARs can help identify functional regulatory variants that influence gene expression [43, 44]. We tested eQTL overlap enrichment by comparing each method’s observed overlap count between DAR regions and ROSMAP lead cis-eQTLs with counts from 10,000 method-matched null region sets. ROSMAP consisted of *N*_sample_ = 560 dorsolateral prefrontal cortex tissue samples, and 16,962 lead cis-eQTLs were identified [45]. Multiple DAR sets showed strong evidence of eQTL enrichment (Table 4). Supplementary Table S4 lists the `Consenrich+ROCCO` gene overlaps. `Consenrich+ROCCO` DARs showed the largest eQTL overlap count and largest null-standardized *z*-score. `ROCCO` DARs likewise showed strong evidence of eQTL enrichment, but with a smaller overlap count and lower null-standardized *z*-score than `Consenrich+ROCCO`.

**Table 4:** Enrichment of lead ROSMAP cis-eQTLs in DARs. Differentially accessible chromatin regions (DARs) identified by each method were evaluated for overlap enrichment with lead cis-eQTLs identified by ROSMAP. Null distributions were approximated by calculating overlaps between the lead cis-eQTL set and 10,000 method-matched null region sets generated with `nullranges::bootRanges`. Null-standardized *z*-scores were calculated as (*O* − *µ*_0_)*/σ*_0_, where *O* is the DAR-eQTL overlap count and *µ*_0_ and *σ*_0_ are the mean and standard deviation of the null overlap counts. Empirical upper-tail *p*-values were calculated as (*b* + 1)*/*(*m* + 1), where *b* is the number of null counts at least as large as the DAR-eQTL count and *m* = 10,000.

| DAR Set Method | eQTL Overlaps | Mean Null Overlaps | Null SD | $z$-score | $p$-value |
| --- | --- | --- | --- | --- | --- |
| Consenrich+ROCCO | 149 | 39.0 | 6.47 | 17.0 | $< 1.0 \times 10^{-4}$ |
| ROCCO | 96 | 26.8 | 5.36 | 12.9 | $< 1.0 \times 10^{-4}$ |
| Genrich | 29 | 9.3 | 3.19 | 6.16 | $< 1.0 \times 10^{-4}$ |
| csaw (5 kb background) | 79 | 23.3 | 5.08 | 10.9 | $< 1.0 \times 10^{-4}$ |
| csaw (10 kb background) | 70 | 17.7 | 4.41 | 11.9 | $< 1.0 \times 10^{-4}$ |
| DiffBind (minOverlap=2) | 80 | 27.6 | 5.45 | 9.62 | $< 1.0 \times 10^{-4}$ |
| DiffBind (minOverlap=3) | 78 | 27.2 | 5.43 | 9.36 | $< 1.0 \times 10^{-4}$ |

We also tested overlap enrichment between DARs and ADSP Tier I gene-body loci. The Alzheimer’s Disease Sequencing Project (ADSP) lists a total of *N*_locus_ = 76 Tier I risk loci supported by genome-wide evidence of effects on Alzheimer’s disease risk. In addition to expert review by ADSP’s Gene Verification Committee, these loci were determined according to methodology and standards discussed in [46]. To evaluate the breadth of AD-risk loci encompassed by each method’s DARs, we counted unique ADSP gene-body loci with at least one DAR overlap and compared that count with the same statistic from matched null region sets. Null region sets were designed to control for varying total genomic coverage and peak size distributions across methods (Section 5.6). As shown in Table 5, `Consenrich+ROCCO` had the highest number of overlaps and largest null-standardized *z*-score for ADSP gene-body loci.

**Table 5:** Alzheimer’s Disease Sequencing Project (ADSP) Tier I gene-body locus overlap enrichment for DARs from each method. Observed values report the number of unique ADSP gene-body loci with at least one DAR overlap. Null means, standard deviations, null-standardized *z*-scores, and empirical upper-tail *p*-values were computed from the same statistic over 10,000 method-matched null region sets generated with `nullranges::bootRanges`.

| Method | DARs | DAR bp | ADSP gene-body loci | Null mean | Null SD | $z$-score | $p$-value |
| --- | --- | --- | --- | --- | --- | --- | --- |
| Consenrich+ROCCO | 11,418 | 4,893,150 | 34 | 14.453 | 3.108 | 6.29 | $< 1.0 \times 10^{-4}$ |
| ROCCO | 6,043 | 3,395,950 | 19 | 9.635 | 2.705 | 3.46 | $1.5 \times 10^{-3}$ |
| Genrich | 1,508 | 1,238,482 | 2 | 3.209 | 1.718 | -0.70 | 0.842 |
| csaw (5 kb background) | 3,239 | 2,673,700 | 16 | 5.623 | 2.196 | 4.73 | $< 1.0 \times 10^{-4}$ |
| csaw (10 kb background) | 2,424 | 2,022,250 | 13 | 4.348 | 1.961 | 4.41 | $3.0 \times 10^{-4}$ |
| DiffBind (minOverlap=2) | 5,799 | 3,413,005 | 24 | 9.865 | 2.762 | 5.12 | $< 1.0 \times 10^{-4}$ |
| DiffBind (minOverlap=3) | 5,929 | 3,378,981 | 24 | 10.047 | 2.778 | 5.02 | $< 1.0 \times 10^{-4}$ |

## 3 Discussion

In this work, we addressed the estimation of latent epigenomic regulatory signals in noisy multi-sample datasets. We introduced a novel method, `Consenrich`, that performs this estimation robustly in multiple assays. Importantly, `Consenrich` outputs a genome-wide consensus signal estimate from multiple samples and corresponding model-implied uncertainty. These outputs provide a continuous, cohort-level summary of regulatory landscapes under uncertainty.

Several opportunities for future work remain. First, while the examples evaluated here were focused on bulk tissue data, the state-space model underlying `Consenrich` may be augmented to explicitly account for sparsity in single-cell data [47, 48]. Generally in this scenario, data is aggregated across cells defined to belong to an annotated cell type and peaks called on these “pseudo-bulk” datasets. The number of cells and thus information available for specific cell types may vary widely and with differing noise profiles, making it challenging to provide peak annotations of similar quality across cell types. Another possible future direction involves richer observation models to estimate linear combinations of complementary functional genomics signals. For instance, a combination of accessible chromatin and histone modification signals may reinforce regulatory elements that are only weakly represented in either assay alone, while also helping distinguish accessible but inactive regions from sites with stronger evidence of enhancer or promoter activity. These types of analyses are made possible due to the genome-wide signal estimates produced by `Consenrich`.

## 4 Conclusions

In summary, `Consenrich` provides a practical state-space estimator for multi-sample functional genomics data. In ATAC-seq, DNase-seq, and ChIP-seq examples, `Consenrich` showed strong estimation performance and supported relevant downstream analyses in functional genomics.

## 5 Methods

### 5.1 Genomic Intervals and the State Vector

Let ℒ denote a genomic region, such as a chromosome or chromosome segment, over which the signal of interest is estimated. We partition ℒ into *n* fixed-width genomic intervals ℐ_[1]_, …, ℐ_[*n*]_, each of length *L* base pairs. The default interval length is *L* = 50 bp. Using coordinates relative to the start of ℒ, interval *i* is

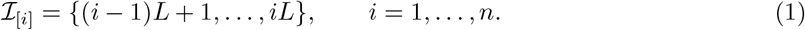

This indexing scheme is illustrated in Figure 6.

**Fig. 6:**
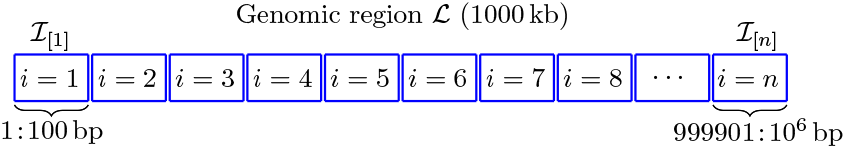
The genomic region ℒ is partitioned into n fixed-width genomic intervals. In this 1000 kb example, each interval ℐ _[i]_ has length L = 100 base pairs, giving n = 10,000 intervals indexed by i = 1, …, n.

The latent signal level at genomic interval ℐ_[*i*]_ is denoted *x*_[*i*]_. To allow the signal to vary smoothly across adjacent intervals, the state also includes a local trend component, denoted 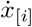. These two quantities define the latent state vector

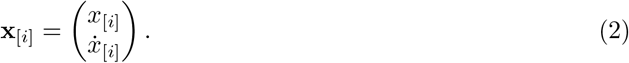

Uncertainty in the latent state is tracked through the corresponding state covariance matrix,

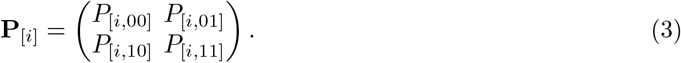

Covariance row and column indices are zero-indexed, i.e., *P*_[*i*,00]_ is the marginal variance of the latent signal level *x*_[*i*]_.

### 5.2 Preprocessing

Let 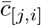 denote the depth-normalized coverage value for sample *j* at genomic interval *i*. By default, sample-level coverage values are transformed into model observations by first normalizing with respect to sequencing depth (effective genome size, scaled to 1*×* genome coverage [49]) and then log-transforming,

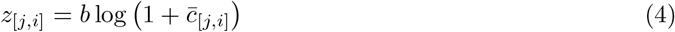

where default 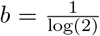. With paired controls, the same log-offset transform is applied to the normalized treatment and control tracks and the transformed control value is subtracted from the transformed treatment value.

### 5.3 Process and Observation Models

At each genomic interval, the fitted state is a weighted combination of a smooth prior prediction and the sample-level observed data: The process model propagates the latent state across genomic intervals, while the observation model specifies how transformed measurements update that prediction. This balance is managed by the *gain*, **K**_[*i*]_ ∈ ℝ^2*×m*^. Informally, the gain is an inverse-shrinkage factor: larger gain entries correspond to observations that are reliable relative to the process model’s predicted state uncertainty, producing a stronger correction toward the data. Conversely, smaller gain entries keep the estimate closer to the prior process prediction (stronger shrinkage to the prior process model).

#### 5.3.1 Process Model

The *process model* specifies how the latent state transitions between genomic intervals during the prior prediction step. Given the forward-filtered state mean and covariance from interval *i* − 1, denoted **x**_[*i*−1|*i*−1]_ and **P**_[*i*−1|*i*−1]_, the linear transition model is

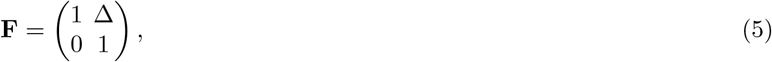

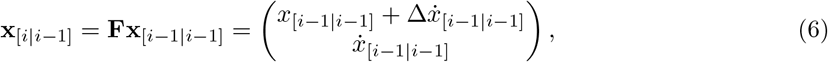

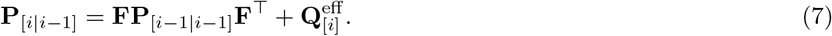

Here Δ *>* 0 controls the propagation of the local trend component across adjacent genomic intervals. In all analyses, we use Δ = 1.

##### Process variance

The process covariance matrix **Q**_[·]_ ≻ 0 reflects varying uncertainty in the smooth prior model: the linear state transition **F** can become unreliable, for example, at genomic intervals where an abrupt shift occurs in the underlying signal. In such instances, the process covariance is adaptively inflated such that estimation favors the observed data more than the prior prediction [50–53].

`Consenrich` uses an *effective process covariance* at each genomic interval,

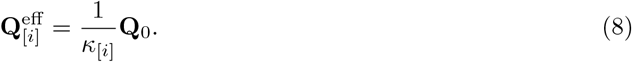

where we refer to *κ*_[*i*]_ as a *process precision multiplier*. Values *κ*_[*i*]_ *<* 1 inflate the effective process covariance, reducing the local influence of the smooth process prediction, while values *κ*_[*i*]_ *>* 1 increase the precision of the model. Note that these precision multipliers are treated as latent variables during estimation and do not require tuning or specification by the user.

Define the process innovation for the transition into interval *i* as

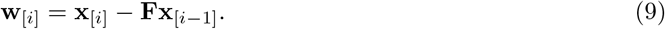

During fitting, this unobserved innovation is summarized through its posterior second moment,

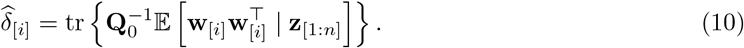

With *ν*_proc_ *>* 0, the precision update is

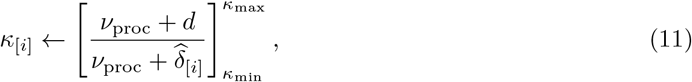

where *d* = 2 is the dimension of the latent state and 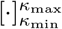 denotes Consenrich’s update-truncation. Large standardized innovations therefore reduce *κ*_[*i*]_, allowing the fitted state to depart from the smooth process prediction at intervals where the transition model is locally unreliable. The fitting procedure described later shows how these precision multipliers are updated together with the smoothed state estimates.

#### 5.3.2 Observation Model

The *observation model* specifies how the transformed sample-level measurements update the prior state prediction *x*[*i*|*i* − 1] at each genomic interval *i*. Measurements are linked directly to the latent signal level of interest, *x*_[*i*]_, the first component of the state vector **x**_[*i*]_. The observation matrix is therefore

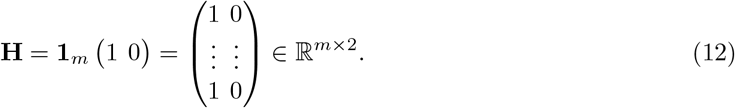

The observation equation is

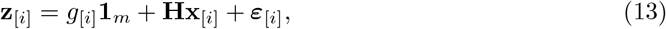

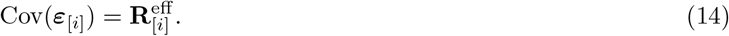

Here *g*_[*i*]_ is a shared background term, and 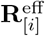 is the effective observation covariance used in the state update. To reduce potential confounding between the shared background and latent signal, the background term is estimated with strong first- and second-order difference penalties,

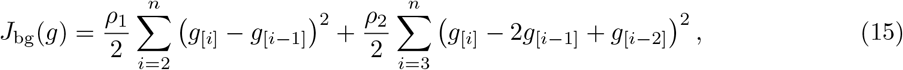

where *ρ*_1_, *ρ*_2_ ≥ 0. In the frequency domain, the combined first- and second-order difference penalty assigns approximate spectral weight 4*ρ*_1_ sin^2^(*ω/*2) + 16*ρ*_2_ sin^4^(*ω/*2). The resulting background update therefore behaves approximately as a low-pass smoother, preferentially assigning broad variation to *g*_[*i*]_ while leaving sharper local variation to the latent signal *x*_[*i*]_. By default, the background penalties are chosen so that an implied dependence span of *g*_[*i*]_ is several multiples larger than the approximate length scale of the latent signal *x*_[*i*]_ (Supplementary Section A.3).

Given the prior state prediction **x**_[*i*|*i*−1]_, define the background-adjusted prediction residuals as

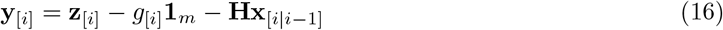

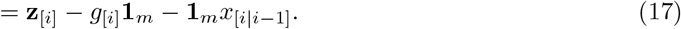

Let *v*_[*j,i*]_ denote the sample-specific observation variance estimated for sample *j* at interval *i* (Section 5.3.2). The effective observation covariance matrices are restricted to be diagonal, following practical conventions in state-space estimation [54], to reduce the number of free variables.

The diagonal entries are

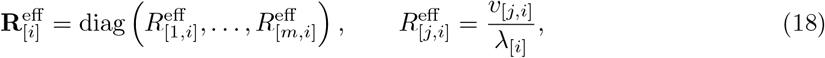

where the multiplier *λ*_[*i*]_ is used for robust interval-level reweighting of the observation variances during fitting (Section 5.4).

The prediction-residual covariance and analytic expression for the gain are

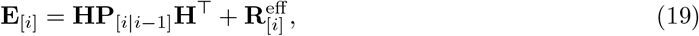

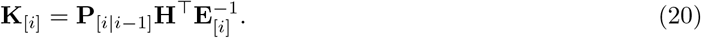

This expression, which is akin to a ratio of the prior state variance to the total residual variance, is consistent with our earlier informal description of the gain as an inverse-shrinkage factor. When the prior state variance is large relative to the observation variance, the gain is large and the state update is more strongly influenced by the data. Conversely, when the prior state variance is small relative to the observation variance, the gain is small and the state update is more strongly influenced by the prior process model.

##### Observation variance

Exact observation covariance matrices are rarely given in practice and must be estimated from the data [55–59]. Further, functional genomics experiments typically contain far fewer independent samples than genomic intervals, making it difficult to estimate sample- and interval-specific observation variances with stability. `Consenrich` therefore shrinks noisy sample- and interval-specific observation variance estimates toward a coarse, replicate-agnostic mean-variance trend fit. Let *L*_[*j,i*]_ denote the local variance estimate for sample *j* at interval *i*, and let 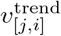 denote the value predicted by the fitted mean-variance trend. The moderated observation variance is then a weighted combination of the local estimate and the trend estimate,

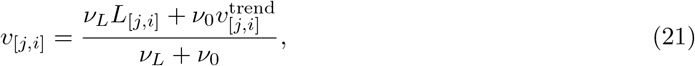

where *ν*_*L*_ is the local evidence weight and *ν*_0_ is the fitted prior strength. This construction resembles empirical Bayes variance shrinkage methods for high-dimensional data [20, 60–62], but is used in our state-space setting with serially dependent data to initialize working observation variances for the Kalman–RTS updates. Figure 7 shows this moderation in an ATAC-seq example, where local per-sample variance curves tighten toward the fitted mean-variance trend while retaining sample-level variance structure. During fitting, the observation variances enter the diagonal effective observation covariance as

**Fig. 7:**
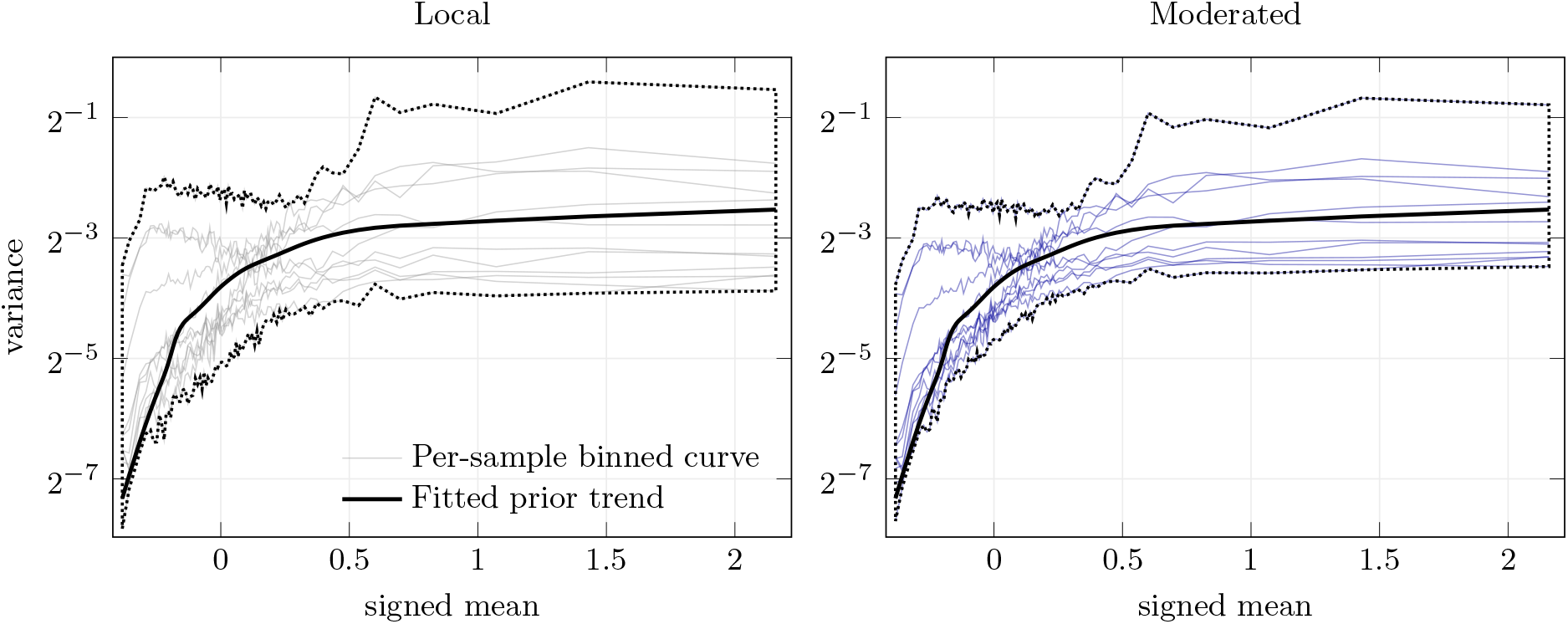
Local vs. moderated observation-variance curves. In the Het10 ATAC-seq dataset, intervals were binned by signed mean, and each thin curve shows the median variance for each sample within each bin. The black curve is the fitted pooled prior trend. Dotted envelopes trace the range spanned by the sample curves and prior trend in each panel. The right panel shows the tighter moderated envelope around the prior trend, indicating global shrinkage toward the prior with retention of sample-level variance structure.

**Fig. 8:**
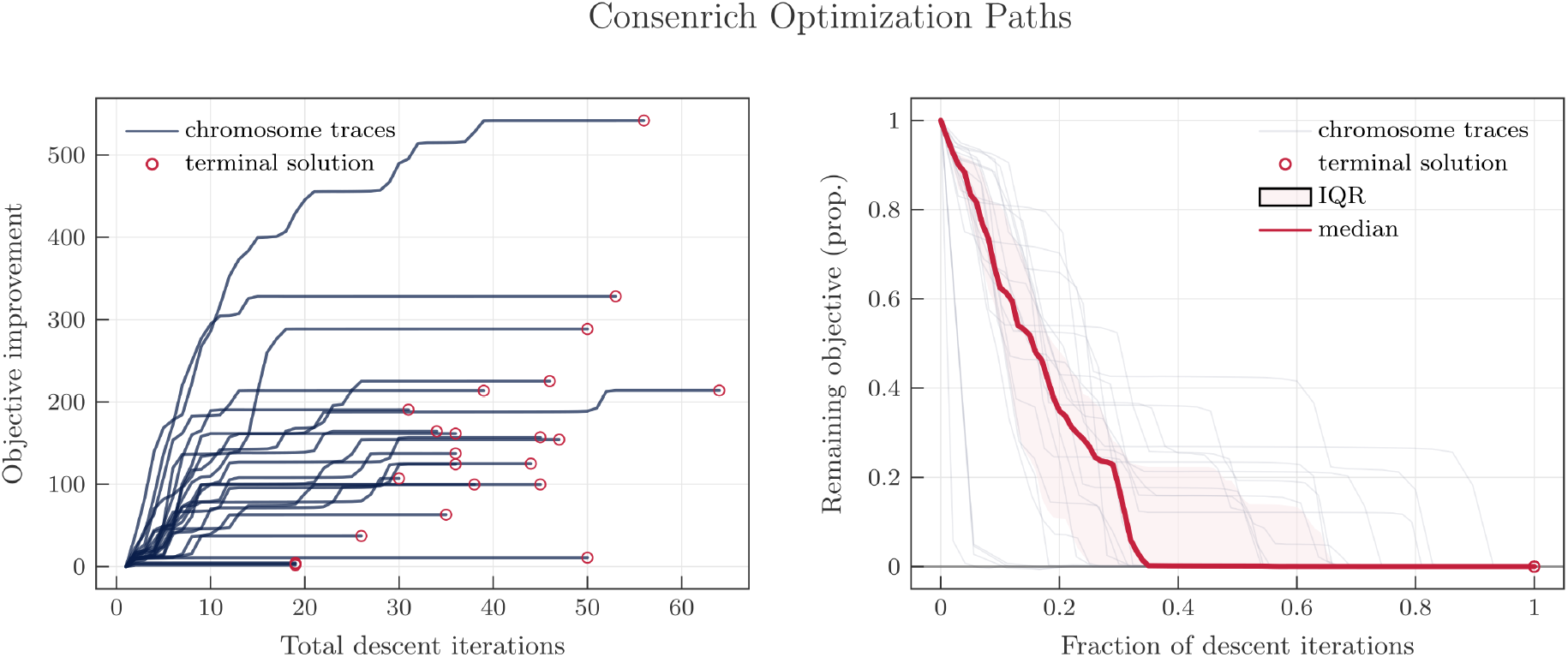
Chromosome-level objective traces during fitting. Empirical convergence trends for each chromosome in the POLR2A ChIP-seq dataset are plotted. In the left panel, cumulative objective improvement is shown against the total number of iterations for each chromosome, with open circles marking terminal chromosome-specific solutions. In the right panel, each trace is rescaled by its descent-iteration span and objective gap, giving the remaining objective proportion along the normalized descent path. The cardinal-red curve gives the pointwise median across chromosomes, and the shaded band gives the interquartile range. The decreases and plateaus near zero agree with the monotonic convergence result in Supplementary Section A.1.

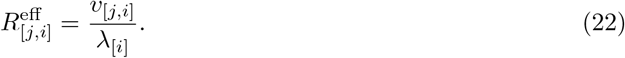

The multiplier *λ*_[*i*]_ is interval-level, so it rescales all samples at interval *i* together, but across-sample variance structure is not modified. See A.2 for complete details.

### 5.4 Model Fitting

Consenrich is fit with a blockwise optimization scheme based on expectation–conditional-maximization (ECM) [63, 64]. First, treating nuisance quantities (background, observation and process precision multipliers) as fixed, the E-step runs the Kalman–RTS smoother and computes posterior moments for the latent states. Then, holding the moments as constant, the nuisance quantities are updated to optimize the conditional objective. The E-step and conditional M-step are repeated until convergence (Section A.1).

Let **z**_[*i*]_ = (*z*_[1,*i*]_, …, *z*_[*m,i*]_)^⊤^ denote the observations at genomic interval *i*. Let

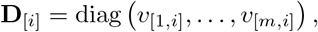

and define

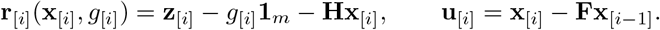

Let **m**_0_ and **P**_0_ ≻ 0 denote, respectively, the fixed mean and covariance of the initial-state prior for **x**_[1]_. The objective function is

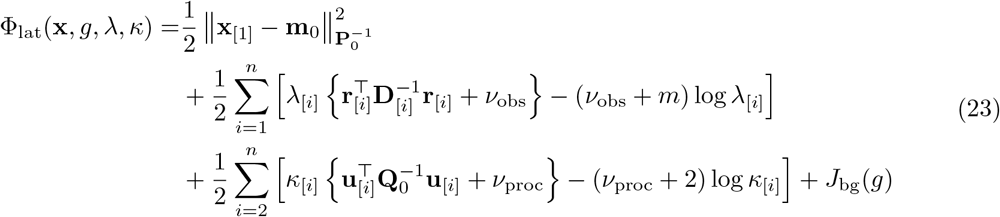

ignoring constants. During fitting, Φ_lat_ is replaced by its conditional average implied by the Kalman– RTS posterior moments from the E-step. The effective observation covariance at interval *i* is

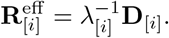

For fixed *g, λ*, and *κ*, the E-step runs a Kalman forward pass followed by a Rauch–Tung–Striebel backward pass, yielding the Kalman–RTS state and covariance estimates

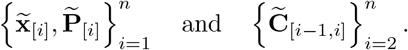

With these fixed, *T*_*λ*_ and *T*_*κ*_ update the observation and process precision multipliers. The observation update uses the Studentized residual term implied by 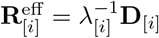, and the process update is given by Equation 11.

For a fixed across-replicate background track, the inner ECM cycle in pass *k* is

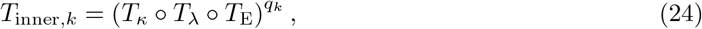

where *T*_E_ denotes the Kalman forward pass followed by the RTS backward pass and *q*_*k*_ is the number of fixed-background cycles.

The across-replicate shared background is then updated by solving the penalized weighted least-squares problem defined by the observation model and Equation 15. Denoting this conditional update by *T*_*g*_, and writing

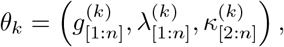

one outer pass is

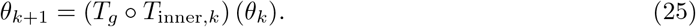

The vector *θ*_*k*_ contains only nuisance quantities. The latent-state posterior moments are recomputed by *T*_E_.

This procedure is outlined in Algorithm 1. For the stopping rule, let 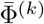 denote the penalized objective per matrix element after outer pass *k*, and let

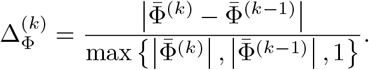

Similarly, define

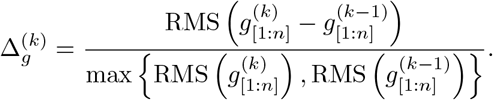

In all analyses, *ε*_Φ_ = 10^−4^, *ε*_*g*_ = 10^−2^, and *k*_min_ = 3. Monotonic convergence is addressed in Supplementary Section A.1.

#### Algorithm 1 Consenrich: Blockwise Fitting

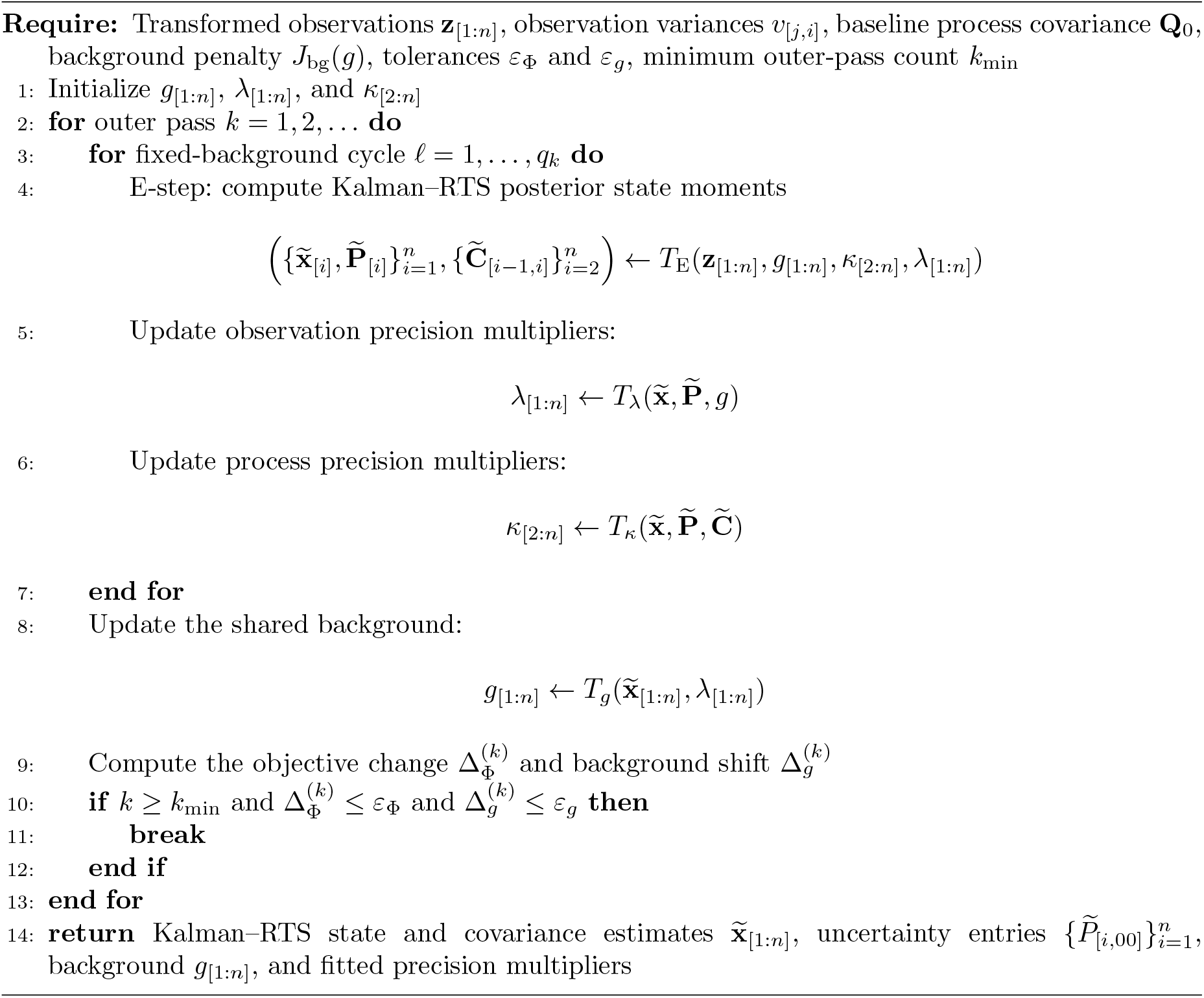

### 5.5 Evaluation datasets

A detailed listing of individual accessions for each evaluation dataset is in Supplementary Section B. Briefly, for signal enrichment analyses, lymphoblastoid ATAC-seq alignment files were downloaded from the ENCODE Portal; GM12878 lymphoblastoid reference annotations were downloaded from ENCODE SCREEN and the Human Heterochromatin Database (HHCDB); POLR2A ChIP-seq treatment and control alignment files from colon tissue samples were downloaded from the ENCODE Portal; and H3K4me1 ChIP-seq alignment files were downloaded from the ENCODE Portal. For differential-analysis comparisons, DNase-seq alignment files from Alzheimer’s disease and nonAD dorsolateral prefrontal cortex donors were downloaded from the ENCODE Portal; ROSMAP lead cis-eQTLs were downloaded from the Synapse/AD Knowledge Portal eQTL meta-analysis resource; and Alzheimer’s disease risk loci were downloaded from ADSP [65].

### 5.6 Synthetic Datasets and Empirical Null Distributions

To generate *Noisy* ATAC-seq samples, we introduced contamination at the sequence alignment level. Unrelated, low-quality alignments were merged with ChIP-seq control alignments and subsampled to generate *m* = 10 total *Noisy* samples with both low signal-to-noise ratios (SNR) and piecewise artifacts capable of distorting true signals (see Supplementary Section B.1 for additional details).

To generate *Null* samples, we constructed empirical null distributions using the segmented block bootstrap implemented in `nullranges::bootRanges` [66]. Here, the null hypothesis is that the observed set of genomic features are exchangeable with features sampled from genomic blocks of the same gene-density state, such that any apparent enrichment in the observed feature set reflects gene-density-matched genomic structure rather than a specific association with the feature being tested. Unlike uniform shuffling, `bootRanges` resamples contiguous genomic blocks, preserving local spatial clustering among nearby features. We used the gene density segmentation to sample within matched gene-density segments. For each replicate, we recomputed the same enrichment statistic on the boot-strapped ranges, yielding an empirical null distribution against which the observed statistic was compared.

### 5.7 Putative Positive

For ChIP-seq assays (H3K4me1, POLR2A), we defined putative target-positive regions from ENCODE cell-line peak calls generated by ENCODE ChIP-seq processing pipelines [67]. Per-experiment peaks with an ENCODE *q*-value score ≥ 3.0 were retained if supported by 5 or more independent experiments. For open chromatin assays, we defined putative accessibility-positive regions from ENCODE cell-line DNase-seq hotspot calls generated by the ENCODE DNase-seq processing pipeline [68]. Peaks with an ENCODE *q*-value score ≥ 3.0 with support from 50 or more independent experiments were retained. Peak support across experimental outputs was calculated using `BEDTools multiinter` [69].

### 5.8 Other Software

For the signal enrichment experiment, the `MACS3 FE (cmbreps)` and `MACS3 logFE (cmbreps)` bench-marks mimic ENCODE’s signal quantification protocols for ATAC-seq and ChIP-seq [9, 23, 24]. `DiffBind` [10] was run on DNase BAMs and matched narrowPeak files from the corresponding ENCODE experiments. Consensus peak sets were formed using `minOverlap` = 2 and `minOverlap` = 3. `csaw` [19] was run directly from the input BAMs using 100 bp windows spaced every 50 bp. Background bin widths of 5 kb and 10 kb were used to normalize the final count matrix. Neighboring windows are summarized with `mergeResults` using `tol` = 500 and `max.width` = 1000.

### 5.9 Differential analyses

With the exception of `csaw`, differential analysis was performed using a multi-step workflow involving `RUVSeq` [70] and `DESeq2` [20]. Donor age was specified as a covariate in the DESeq2 design formula along with *K* = 3 latent sources of variation from a reduced-rank representation of count data at negative control regions (`RUVSeq::RUVg`) [70]. The following design formula was supplied to DESeq2:

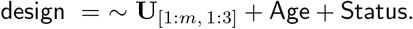

An absolute fold-change threshold log_2_ |FC| *>* 0.25 and FDR-corrected *p*_adj_ *<* 0.05 were specified to determine significant disease-specific changes in chromatin state.

### 5.10 Pathway enrichment analyses

`clusterProfiler::enrichGO` was run with a false discovery rate (FDR)-adjusted *p*-value thresh-old (`p.adjust < 0.01`) and semantic similarity cutoff (`clusterProfiler::simplify(cutoff=0.70)`) [71] to determine significantly enriched, nonredundant Gene Ontology (GO): Biological Process terms related to the differentially accessible regions.

### 5.11 DAR-eQTL and ADSP locus overlaps

To test for enrichment of disease-associated differentially accessible regions (DARs) among lead cis-eQTLs from the ROSMAP meta-analysis, we used 10,000 null sets of genomic regions generated by `bootRanges` as described in Section 5.6. The observed overlap count was compared against the empirical null distribution to obtain a *p*-value for enrichment. For the ADSP locus test, the observed statistic was the number of unique ADSP Tier I gene-body loci with at least one DAR overlap. Multiple DARs hitting the same ADSP locus were counted once, and the same statistic was recomputed for each matched null set.

## Declarations

### Availability of Data and Materials

The sequence alignment files used as input to evaluate `Consenrich` are publicly available from ENCODE [2]. Full details and accession codes can be found in Supplementary Sections B.1, B.2, and B.3. A complete software implementation of `Consenrich` is available at https://github.com/nolan-h-hamilton/Consenrich under the MIT license.

### Author Contributions

**NHH**: Conceptualization, Methodology, Formal analysis, Software, Validation, Writing – original draft, Writing – review and editing. **YCH**: Data Curation, Formal analysis, Experiment design, Validation, Writing – original draft, Writing – review and editing. **BDM**: Data Curation, Experiment Design, Validation, Writing – review and editing. **MIL**: Experiment design, Supervision, Writing – review and editing. **TSF**: Experiment design, Funding acquisition, Resources, Supervision, Writing – review and editing.

